# TraRECo: A Greedy Approach based *de novo* Transcriptome Assembler with Read Error Correction using Consensus Matrix

**DOI:** 10.1101/216077

**Authors:** Seokhyun Yoon, Daeseung Kim, Keunsoo Kang, Woong June Park

**Affiliations:** Department of Electronics Eng., College of Engineering, Dankook University, Yongin-si, Korea; Department of Microbiology, College of Natural Sciences, Dankook University, Cheonan-si, Korea; Department of Molecular Biology, College of Natural Sciences, Dankook University, Cheonan-si, Korea

**Keywords:** RNA-Seq, *de novo* Transcriptome assembly, greedy approach, consensus matrix, read error correction

## Abstract

**Background:** Challenges in developing a good *de novo* transcriptome assembler include how to deal with read errors and sequence repeats. Almost all *de novo* assemblers utilize de Bruijn graph, which has a complexity linearly growing with data size while suffers from errors and repeat. Although one can correct errors by inspecting topological structure of the graph, it is an uneasy task when there are too many branches. There are two research directions: improving either graph reliability or path search precision. We focused on improving the reliability.

**Results:** We present TraRECo, a greedy approach to *de novo* assembly employing error-aware graph construction. The idea is similar to overlap-layout-consensus approach used for genome assembly, but is different in that consensus is made through the entire graph construction step. Basically, we built contigs by direct read alignment within a distance margin and performed junction search to construct splicing graphs. While doing so, however, a contig of length *l* was represented by 4×1 matrix (called consensus matrix), of which each element was the base count of aligned reads so far. A representative sequence is obtained, by taking majority in each column of the consensus matrix, to be used for further read alignment. Once splicing graphs were obtained, we used IsoLasso to find paths with noticeable read depth. The experiments using real and simulated reads showed that the method provides considerable improvements in sensitivity and reasonably better performances when comparing both sensitivity and precision. This could be achieved by making more erroneous reads to be participated in graph construction, which, in turn, improved the depth information quality used for the subsequent path search step. The results for simulated reads showed also challenges are still remaining since non-negligible percentage of transcripts with high abundance were not recovered by the assemblers we considered.

**Conclusion:** *de novo* assembly is mainly to explore not-yet-discovered isoforms and must be able to represent as much reads as possible in an efficient way. In this sense, TraRECo provides us a potential alternative to improve graph reliability, even though the computational burden can be much higher than single *k*-mer de Bruijn graph approach.

## Background

Low-cost and high-throughput transcriptome profiling (RNA-seq) based on next generation sequencing technology ignited recent development of many software tools for the transcriptome assembly to explore novel isoforms, co-expression of related genes, differentially expressed genes, and so on. The transcriptome assembly to analyze such RNA-seq data are categorized into two: one is reference-based assembly and the other is *de novo* assembly. In the reference-based assembly, such as cufflinks (Trapnell, *et. al.* 2010), Scripture (Guttman, *et. al.* 2010), IsoLasso (Li, *et. al.,* 2011), and Isoinfer (Feng, *et. al.,* 2010), raw RNA-seq reads are first mapped to a reference genome, for example by using a splice-aware aligner such as TopHat (Trapnell, *et. al.* 2009) and use gene-annotation information to explore expressed transcript/isoforms and their expression level. While, in *de novo* assembly, raw RNA-seq data are aligned with each other to construct directly a graph representing possible splicing patterns, by which isoform detection and expression-level estimation can be performed. Although reference-based assembly provides more accurate isoform detection and abundance estimation, it highly relies on prior knowledge of known genes, i.e., it requires any known genome and annotation, and can neither be used for finding novel isoforms, including fusion genes, nor applied to the species without reference genome. Due to these reasons, recent researches are focusing more on *de novo* assembly and many +*novo* assemblers have been proposed so far, including Trans-ABySS (Robertson, *et. al.* 2010), Trinity (Grabherr *et al.*, 2011), Velvet-OASES (Zerbino *et al.*, 2008, Schulz *et al.*, 2012), IDBA-tran (Peng *et al.*, 2013), SOAPdenovo-Trans (Xie *et al.,* 2014) and, more recently, Bridger (Chang *et al.*, 2015), and BinPacker (Liu *et al.*, 2016).

Challenges to develop a good transcriptome assembler include how to deal with read errors and sequence repeats that frequently occur in eukaryotes. Almost all the *de novo* transcriptome assemblers utilize de Bruijn graph, which is constructed by converting base sequences into *k*-mer sequences and counting *k*-mers. This approach was widely adopted by most of the *de novo* transcriptome assemblers since its complexity grows linearly with the read data size. A weakness of de Bruijn graph-based approach, however, is the difficulty in correcting read errors. One way to deal with read errors is pre-error correction, such as Trimmomatic (Bolger *et al.*, 2014), and QuorUM (Marais *et. al.,* 2015) based on the quality score of reads, which, however, is a little bit risky as the quality score tells us only probability. Other pre-error correction algorithms include Coral (Salmela *et* al., 2011), SEECER (Le *et* al., 2013) and Rcorrector (Song *et* al., 2015), some based on multiple alignments and some on Probabilistic approach. Another way to handle read error, especially in de Bruijn graph-based approach, is to utilize topological structure of the graph, i.e., read errors make many branches with relatively low read coverage and one can simply remove those suspicious branches or merge them to the main branch if it is possible to identify the main one. Another approach has been recently proposed in IDBA-tran where the authors employed an iterative remapping of reads to the de Bruijn graph with increasing *k*-mer values. Another drawback of de Bruijn graph based approach is that it cannot utilized full connection information of a read. Most of the de Bruijn graph-based *de novo* assemblers use *k* around 30, which is much smaller than the typical read length of 75 or 100. This means that the effective read length is only 30. As a matter of fact, with small value of *k,* two or more isoforms that have common sequences of length longer than *k* can be merged into a single (sometimes huge) graph, making isoform detection complicated and decreasing the prediction accuracy. To overcome this tradeoff, (Schulz *et al.*, 2012) proposed combining of splicing graph obtained with different*k*-mers, which was shown to be effective to resolve such problem.

Based on the researches in *de novo* transcriptome assembly so far, there are two main research directions. One is to improve precision in plausible path search for given splicing graphs as in Bridger (Chang *et al.*, 2015) and BinPacker (Liu *et al.*, 2016) and the other is to improve reliability in splicing graph construction, for example by utilizing multi *k*-mers approach as in OASES (Schulz *et al.*, 2012) and IDBA-tran (Peng *et al.*, 2013). In this study, we took the latter direction based on a greedy approach employing error-aware construction of splicing graph.

To combat read errors and sequence repeat, we built splicing graphs by directly aligning reads, as did in the conventional genome assembly, which made us resort full connection information of a read even though the running time can be much longer than the single *k*-mer de Bruijn graph-based approach. As provided in (Li, *et al*., 2012), there are two approaches to genome assembly: one is overlap-layout-consensus (OLC) and the other de Bruijn graph approach. The former first gathers alignment information between all pairs of reads to build overlap graph, bundle stretches to obtain contigs and finally align contigs to correct possible errors by taking *consensus* at each position of aligned contigs. The proposed approach is similar to the former, but with clear difference in that aligning the reads and taking consensus (error correction) are performed simultaneously in the course of entire graph construction step. In our scheme, a contig of length *l* was represented by a consensus matrix of size 4xl, of which each element is the base count of the reads aligned to this contig, and a representative sequence, of which each letter corresponds to the majority (the row index with the highest count) at each column, is used for further read alignments. Once overlap detected for a read within some distance margin, we update its consensus matrix by increasing those elements corresponding to each letter of the read. As the result, the consensus matrix tracks alignment records throughout the entire assembly process and can be used later for joint isoform detection and abundance estimation. The procedure can improve the reliability of the splicing graph by making more erroneous reads to be participated in the graph construction, which, in turn, improves the quality of read coverage depth used for the subsequent plausible path search step. Although two or more isoforms having similar sequences could be merged to a single graph, it was not mostly due to short repeats, but to similar sequences as we could set minimum overlap threshold much longer than that used in the de Bruijn graph-based approach. As will be discussed in detail later, one can check if similar sequences were merged or not by inspecting whether any other element except the representative was outstanding or conspicuous in each column.

## Results

To demonstrate the effectiveness of the proposed scheme, we used three data sets: one was simulated reads with the exact set of target references and their expression levels and others were real RNA-seq data, one for human and the other for mouse, from gene expression omnibus available at (https://www.ncbi.nlm.nih.gov/geo/).

*Using BLASTN*: The primary criterion for performance comparison was the number of ‘distinct’ pair of reference-candidate transcript for given minimum target coverage, where the coverage was the ratio of the alignment length covered by a candidate (assembled) transcript to the length of the reference transcript. The coverage can be obtained by using BLASTN software (Camacho et. al., 2009) which align query sequences to subject sequence database. We used reference transcripts as a query while candidates as subject sequence database. By running BLASTN, we obtain the following for each query (reference) - subject (candidate) sequence pair: *L*_*q*_: query sequence length,*L*_*s*_: subject sequence length and *L*_*a*_: alignment length. The coverage is defined simply as the maximum of *L*_*a*_/*L*_*q*_ over all candidates. We set the maximum number of target sequence option to 1, which means we selected the best match candidate for each reference transcript. Note that, even with this option, a candidate transcript can be matched to a multiple references. Hence, we looked up the match list to select only one reference-candidate pair for each candidate transcript.

*Sensitivity and precision as primary criteria*: If the exact set of transcripts that reside in a sample is known, one can measure the sensitivity, the percentage of the recovered transcripts in the total number of target references, which we could provide for the simulated read. For real reads, however, one do not know the exact set of targets. So, for real data, we compared the number of recovered transcript. In the evaluation of sensitivity, we basically allow each candidate transcript and reference can have only one match. Sometimes, however, a multiple of references are matched to one candidate transcript due to artificial gene fusion. In this case, one can define wide sense (or extended) sensitivity where we allowed multiple transcripts to match with the same candidate for given target coverage. Another primary performance criterion is the precision as a measure of compactness of the assembly. It is defined as the percentage of true positives among all candidate (assembled) transcripts found with a specific assembler. As a matter of fact, there exists trade-off between the two performance criteria (sensitivity and precision) and we need to compare both measures at the same time for a specific target coverage, for which we plotted sensitivity versus precision as shown later.

### 1. Results for Real Reads

#### Data sets

First, we performed *de novo* assembly with two real data sets, one for human and the other for mouse. The human sample was obtained from NCBI website, accession code SRR445718, which was sequenced from embryonic stem cells derived from human pre-implantation embryos and available in the sequence read archives (SRA) format from NCBI. SRR445718 contains approximately 33,000,000 single-ended reads of nominal read length 100 each. The mouse sample was the one tested in many studies on *de novo* assemblers, i.e., accession code SRX062280, and contains approximately 53,000,000 pair-ended reads of nominal read length 76 each.

#### Parameter setting and pre/post processing

We compared TraRECo with some popular *de novo* assemblers, such as Trinity (version 2.4.0), Velvet (version 1.2.01) + Oases (version 0.2.02) and SOAPdenovo-Trans (version 1.01), TransABySS (version 1.5.2), Bridger (version 2014-12-01) and BinPacker (version 1.0). For SOAPdenovo-Trans and Trans-ABySS, we set the *k* value to 31 and 32, respectively, while for Trinity, to 25, the default value. For Bridger and BinPacker, we used *k*-mer length of 25 for human and 31 for mouse as it was suggested in their original paper. With Velvet+Oases, we performed multi-*k* assembly with *k* from 21 to 37 with a step of 4. Parameters for TraRECo was *D*_*th*_ = 0.03~0.06, *O*_*th*_ = 52 for human sample and 44 for mouse (roughly 60% of the read length), *C*_*th*_ = *J*_*th*_ = 24, the default values. The same as other assemblers, we set *L*_*th*_ = 200, i.e., we discarded those candidates of length shorter than 200. As for preprocessing, we first used Cutadapt (Martin 2011) to remove any adaptor sequence remaining in the reads, except for Trinity, for which we enabled Trimmomatic read trimming option, instead of applying Cutadapt. Using the trimmed data, we obtained assembled transcript candidates and, finally, BLASTN was used to align each candidate to reference transcriptome, for which we used Ensembl transcript for human (hg19) and mouse (mm9), respectively, and to finally obtain the number of recovered reference transcripts for given target coverage.

#### Impact of distance threshold (D_th_) and coverage depth threshold (CD_th_) in TraRECo

Table 1 and Fig.1 shows the results for SRR445718 while Table 2 and Fig.2 for SRX062280. Fig.1 and 2 show precision versus the number of recovered transcripts for target coverage (recovered percentage) of 95% (a) and 80% (b). In Fig.1 and 2, the results for assemblers other than TraRECo are shown as a single point, while the results for TraRECo is shown as a curve, where each point in the curve corresponds to *CD*_*th*_ of 0, 1, 2, 4, 8, 12 and 16, respectively. Note that TraRECo detects isoforms and estimates their abundances (expression level) jointly and *CD*_*th*_ was the abundance (coverage depth) cutoff with which we discarded those candidates with their abundance estimate below this value, as most of them are highly likely to be artifacts of read errors. *(CDth* = 0 means that we consider all the paths obtained from the final splicing graph regardless of the abundance estimates.) With various values of this cutoff, the performance of TraRECo is shown as a line for given *D*_*th*_. To show the impact of distance threshold, we ran TraRECo with *D*_*th*_ = 0.04, 0.05 and 0.06, respectively for human and 0.03, 0.04 and 0.06, respectively for mouse, even though Table 1 shows only for *D*_*th*_ = 0.06. The performance of TraRECo in Fig.1 and 2 show the impact of *D*_th_ on the performance of TraRECo, where one can observe the improvements both in precision and sensitivity with larger distance threshold up to *D*_*th*_ = 0.06, after which, however, no further improvement in sensitivity was observed, while the precision was slightly improved. From the performance curves for TraRECo, one can clearly see the performance improvement with higher value of *D*_*th*_, up to 0.06, which shows the validity of the read error correction using consensus matrix at least to some extent. The results in this paper suggests that *D*_*th*_ = 0.06 was reasonable choice to obtain a good result for real data, even though it must be chosen carefully according to error statistics as suggested from the results for simulated read to be shown later.

**Table 1.** A comparison of the number of transcripts found and matched to a reference for the single-ended reads SRR445718 (Human sample). Target coverage = 95%, 90% and 80%. For TraRECo, the results with *D*_*th*_ =0.06 and *CD*_*th*_ =0, 1, 2, 4, 6, 8, 12 and 16 are shown.

| Assembler | # of<br>candidates<br>found | # of transcripts matched<br>for Target coverage of |  |  |
| --- | --- | --- | --- | --- |
| | | $\geq 95\%$ | $\geq 90\%$ | $\geq 80\%$ |
| SOAPdenovo-trans | 48,462 | 2,613 | 3,522 | 5,349 |
| Trans-ABYSS | 87,686 | 3,626 | 4,781 | 6,967 |
| Trinity | 82,865 | 4,008 | 5,279 | 7,680 |
| BinPacker | 20,612 | 3,486 | 4,314 | 5,685 |
| Multi-k OASES | 202,152 | 5,120 | 6,737 | 9,520 |
| Bridger | 58,217 | 4,013 | 5,093 | 7,017 |
| TraRECo, $D_{th}=0.06$, $CD_{th}=0$ | 219,402 | 5,366 | 7,010 | 10,168 |
| TraRECo, $D_{th}=0.06$, $CD_{th}=1$ | 118,342 | 4,845 | 6,308 | 9,078 |
| TraRECo, $D_{th}=0.06$, $CD_{th}=2$ | 109,551 | 4,774 | 6,209 | 8,901 |
| TraRECo, $D_{th}=0.06$, $CD_{th}=4$ | 79,073 | 4,537 | 5,890 | 8,393 |
| TraRECo, $D_{th}=0.06$, $CD_{th}=6$ | 58,396 | 4,217 | 5,436 | 7,642 |
| TraRECo, $D_{th}=0.06$, $CD_{th}=8$ | 47,095 | 3,937 | 5,026 | 6,973 |
| TraRECo, $D_{th}=0.06$, $CD_{th}=12$ | 35,506 | 3,396 | 4,305 | 5,880 |
| TraRECo, $D_{th}=0.06$, $CD_{th}=16$ | 28,997 | 2,959 | 3,749 | 5,089 |

\* Except for Trinity, Cutadapt was used for read data trimming.

**Fig.1.**
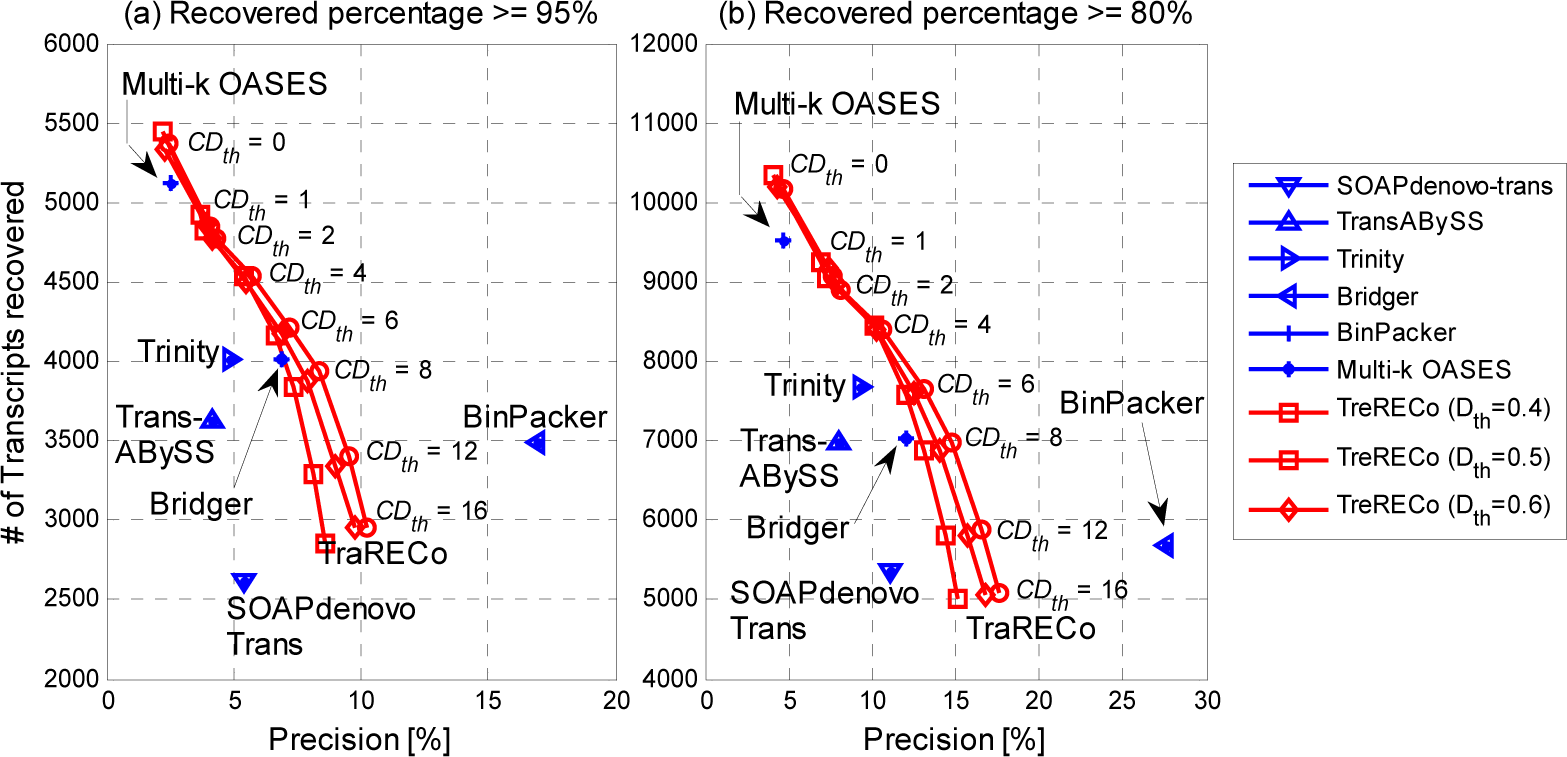
The number of transcripts recovered versus precision for the single-ended read data SRR445718 (Human sample). Target coverage = 95% (a) and 80% (b), respectively.

**Table 2.** A comparison of the number of transcripts found and matched to a reference for the pair-ended read data SRX062280 (Mouse sample). Target coverage = 95%, 90% and 80%. For TraRECo, the results with *D*_*th*_ =0.06 and *CD*_*th*_ =0, 1, 2, 4, 6, 8, 12 and 16 are shown.

| Assembler | # of<br>candidates<br>found | # of transcripts matched<br>for Target coverage of |  |  |
| --- | --- | --- | --- | --- |
| | | $\geq 95\%$ | $\geq 90\%$ | $\geq 80\%$ |
| SOAPdenovo-trans | 48,234 | 4,902 | 6,088 | 7,634 |
| Trans-ABYSS | 70,052 | 8,202 | 9,673 | 11,680 |
| Trinity | 71,415 | 10,247 | 11,645 | 13,543 |
| BinPacker | 34,838 | 8,979 | 10,151 | 11,685 |
| Multi-k OASES | 174,724 | 11,256 | 13,224 | 15,789 |
| TraRECo, $D_{th}=0.06$, $CD_{th}=0$ | 240,907 | 11,864 | 13,980 | 17,076 |
| TraRECo, $D_{th}=0.06$, $CD_{th}=1$ | 82,522 | 10,480 | 12,355 | 14,927 |
| TraRECo, $D_{th}=0.06$, $CD_{th}=2$ | 78,226 | 10,347 | 12,183 | 14,688 |
| TraRECo, $D_{th}=0.06$, $CD_{th}=4$ | 65,823 | 9,978 | 11,706 | 14,080 |
| TraRECo, $D_{th}=0.06$, $CD_{th}=6$ | 52,465 | 9,555 | 11,166 | 13,353 |
| TraRECo, $D_{th}=0.06$, $CD_{th}=8$ | 43,550 | 9,146 | 10,638 | 12,643 |
| TraRECo, $D_{th}=0.06$, $CD_{th}=12$ | 33,704 | 8,313 | 9,614 | 11,282 |
| TraRECo, $D_{th}=0.06$, $CD_{th}=16$ | 28,132 | 7,577 | 8,708 | 10,087 |

\* Except for Trinity, the cutadapt SW was used for read data trimming.

**Fig.2.**
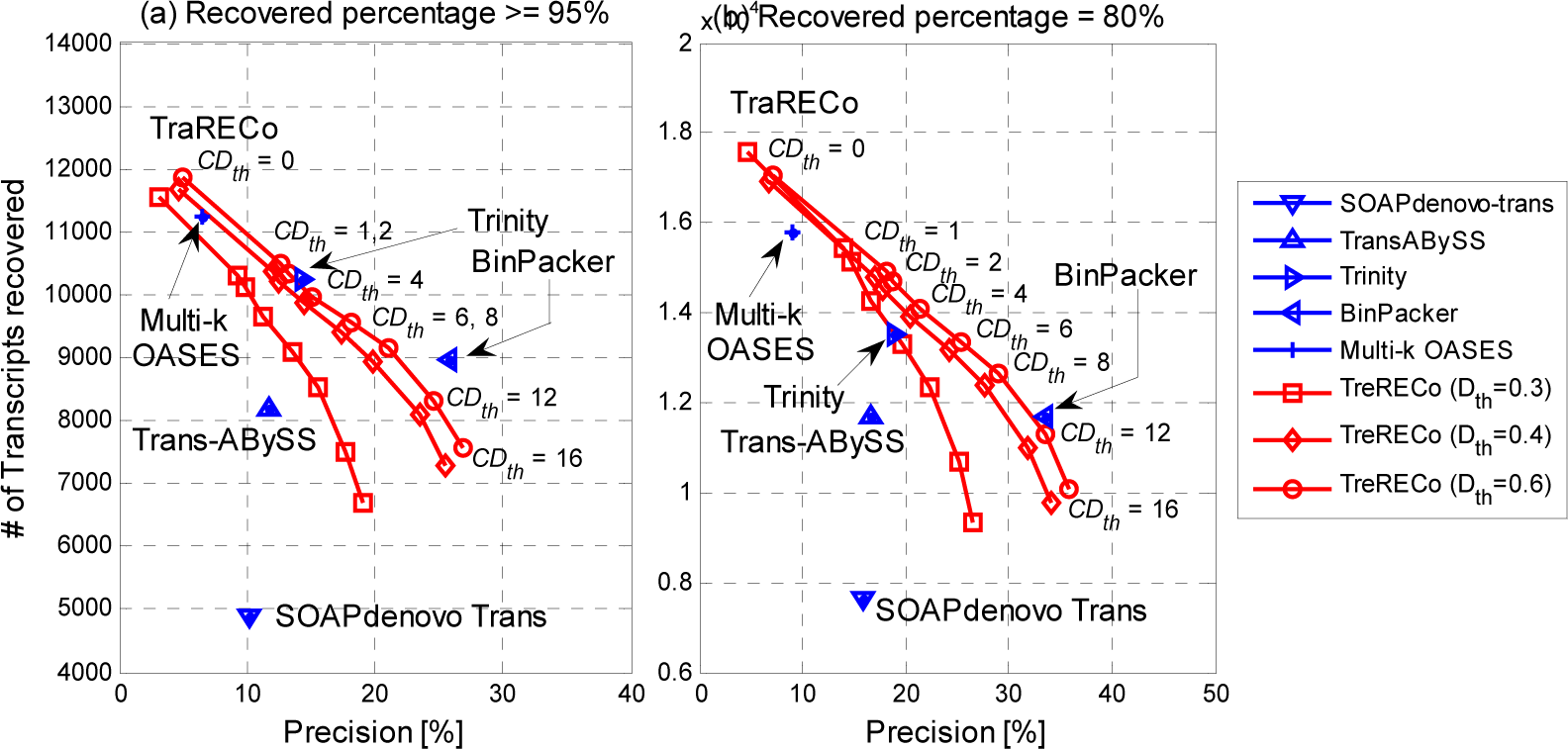
The number of transcripts recovered versus precision for the pair-ended read data SRX062280 (Mouse sample). Target coverage = 95% (a) and 80% (b), respectively.

#### Sensitivity versus precision

In Fig. 1 for human sample, TraRECo showed a better result than most of the assemblers, except for BinPacker, which showed an exceptionally good precision. When considering only the number of transcripts recovered, TraRECo with *D*_*th*_ = 0 and multi-k OASES showed the best results as they found more than 5000 transcripts for 95% target and 9500 for 80%, while most of other assembler found only around 4000 or less for 95% and 7700 or less for 80%. Of course, such high sensitivity in TraRECo with *D*_*th*_ = 0 and multi-*k* OASES might have been obtained only at the cost of precision. For mouse sample shown in Fig.2, when compared at the same precision or at the same sensitivity, TraRECo showed slightly better results than most assemblers, except for Trinity and BinPacker, which showed a better performance than what TraRECo can provides. For mouse sample, we could not obtain the results for Bridger due to runtime errors we couldn’t correct.

#### Transcript length unbalances

In the sensitivity measures obtained here, we allowed only one reference transcript (with the highest alignment length) can be paired with each candidates. However, comparing the length of candidate transcripts with those of their paired reference, one can find that there are big differences between the two. Fig.3 and 4 show scatter plots of the lengths of the candidate (assembled) transcripts and their paired reference for SRR445718 and SRX062280, respectively, where we showed only those for Trans-ABySS, multi-k OASES and TraTECo (*D*_*th*_ = 0.06, *CD*_*th*_ = 4). For human sample, the R^2^ measure for Trans-ABySS, multi-*k* OASES and TraRECo were 0.467 (best), 0. 158 (worst) and 0.398, respectively, and for mouse sample, they were 0.6, 0.324 (worst) and 0.79 (best), respectively. For other assemblers, the R^2^ measure for human (mouse) were 0.382 (0.593), 0.408 (0.511). 0.351 (0.44) and 0.349 (not available), respectively, for SOAPdenovo-trans, Trinity, BinPacker and Bridger.

**Fig.3.**
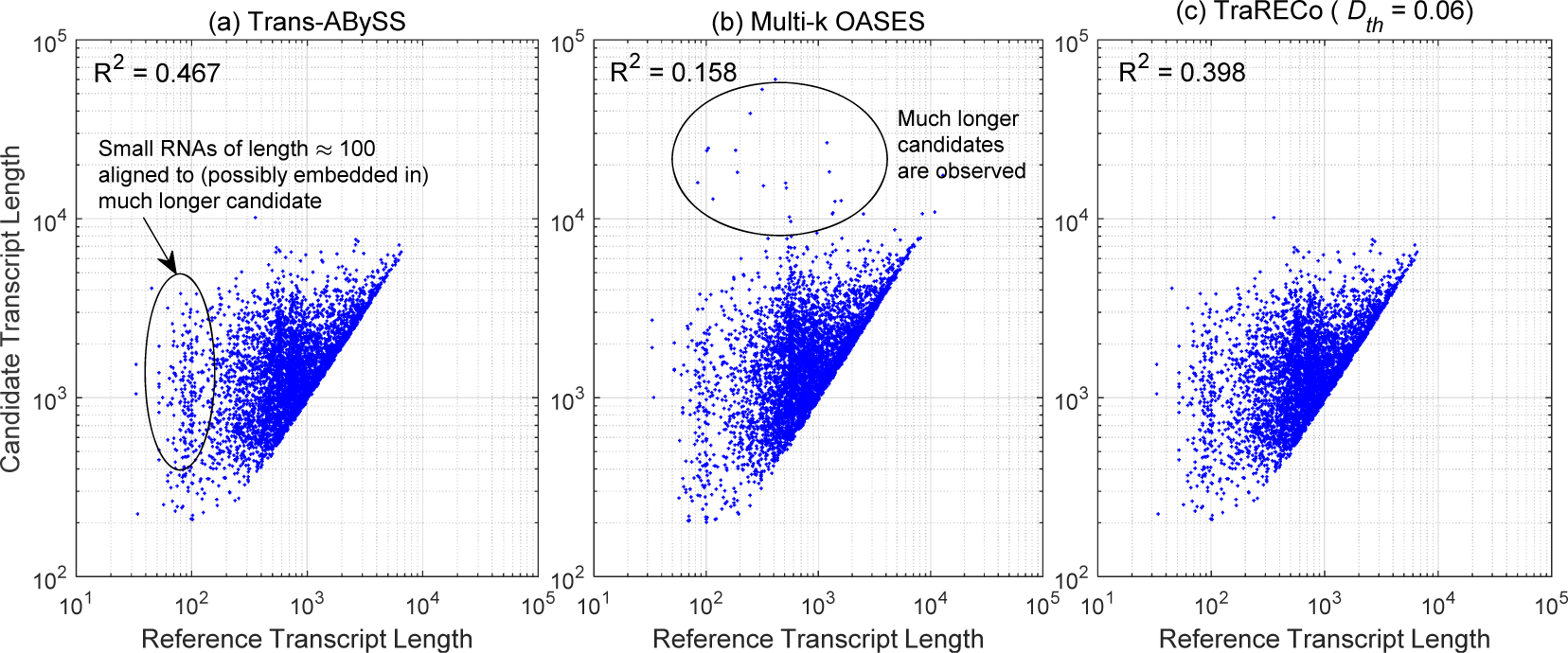
Scatter plot representing length correlation between reference and candidate transcripts matched 13 with 95 or higher coverage of reference for human sample (SRR445718). (a) Trans-ABySS, (b) multi-*k* OASES, (c) TraRECo (*D*_*th*_ = 0.06).

**Fig.4.**
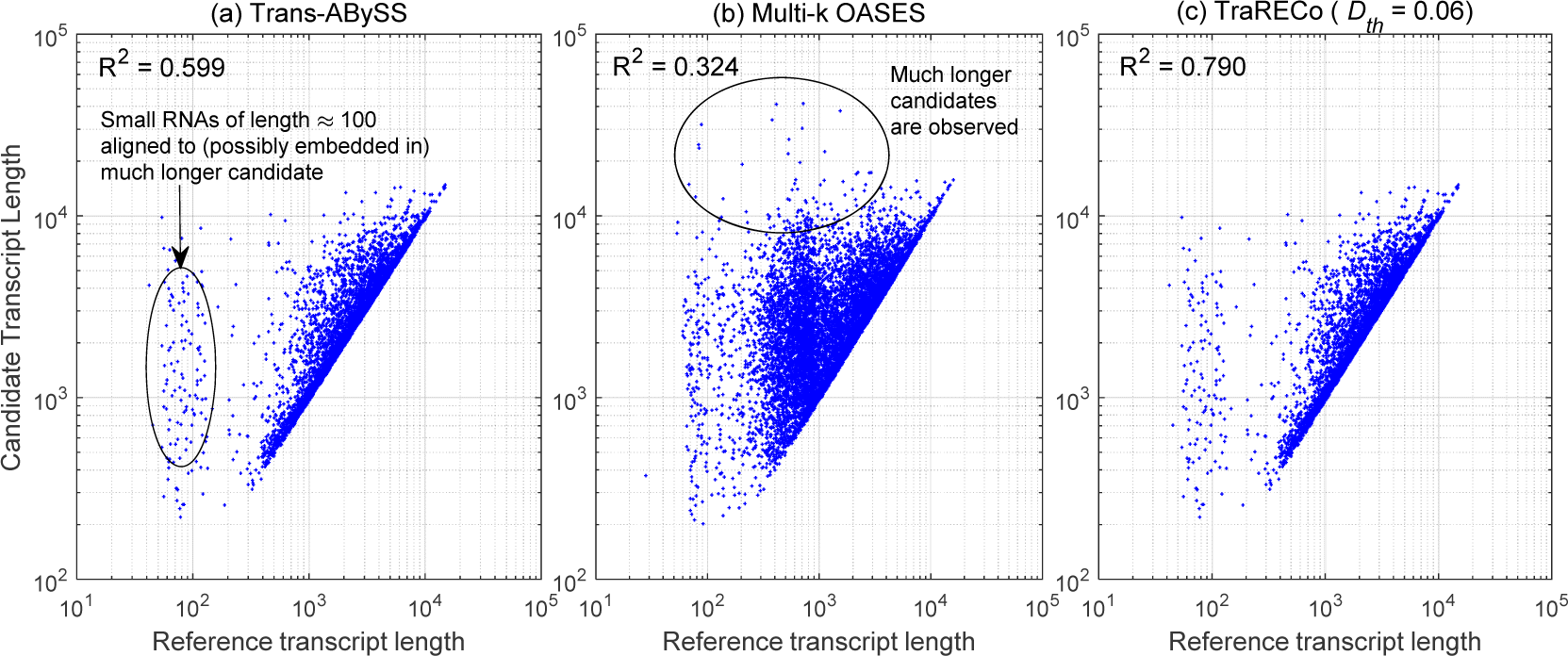
Scatter plot representing length correlation between reference and candidate transcripts matched with 95 or higher coverage of reference for mouse sample (SRX062280). (a) Trans-ABySS, (b) multi-*k* OASES, (c) TraRECo (*D*_*th*_ = 0.06).

#### Wide sense sensitivity (WS sensitivity)

Fig.3 and 4 show that many transcripts, including small RNAs with its length around 100, were merged to a much longer transcripts so that a candidate may represent multiple reference transcripts. Nevertheless, the unbalance between the two lengths does not mean worse performance since they simply stem from artificial gene fusion due to sequence repeat and we could find that there are significant unbalances in all assemblers. Maybe, those small RNAs recovered with *de novo* assemblers were not expressed at all while they were detected because other transcripts contain the same sequences as a part of their entire sequence. Once we consider the artificial gene fusion and the unbalance as a general phenomenon (even though a good assembler should be able to combat sequence repeat in an efficient way), one can check if multiple reference transcripts were merged to one candidate. To take such fusion into account, we define the wide sense sensitivity as the number of reference transcripts that are recovered by any candidate for given minimum target coverage, i.e., by allowing multiple references paired with one candidate. Fig.5 and 6 compare the wide sense sensitivity (the number of references recovered by any candidate) with the (strict sense) sensitivity, which is the one allowing only one candidate for each reference as did before. As shown in Fig.5 and 6, there are noticeable differences between the (strict sense) sensitivity and wide sense sensitivity in all assemblers. Specifically for human sample, wide sense sensitivities were higher than twice of (strict sense) sensitivities for all the assemblers considered, where the largest difference in percent was that of Bridger. The overall wide sense sensitivity looks have similar pattern over those assemblers considered here, while Bridger and Trinity shows quite good performances, for human and mouse, respectively, when considering their precisions shown in Fig. 1 and 2 as well. One thing to note here, however, when considering wide sense sensitivity, one may need to re-define the precision since many candidate transcripts (isoforms) found with each assembler share the same exons and it might be more appropriate to use, for example, the number of nucleotides in the splicing graphs, which are not available for most of the assemblers.

**Fig.5.**
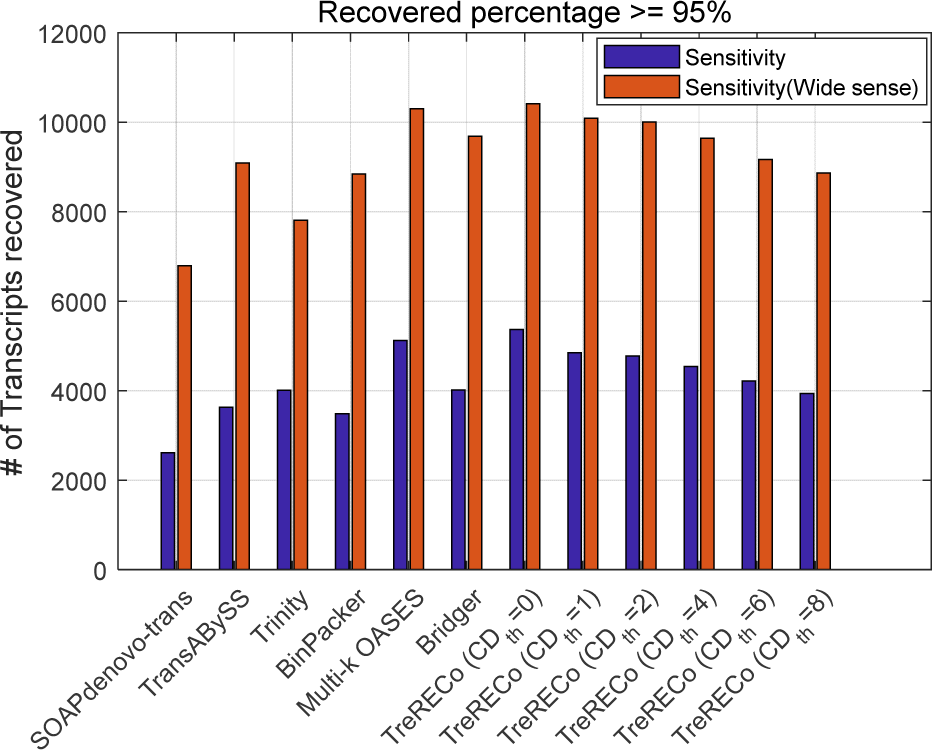
A comparison of (strict sense) sensitivities with wide sense sensitivities for human sample (SRR445718). Target coverage ≥ 95%.

**Fig.6.**
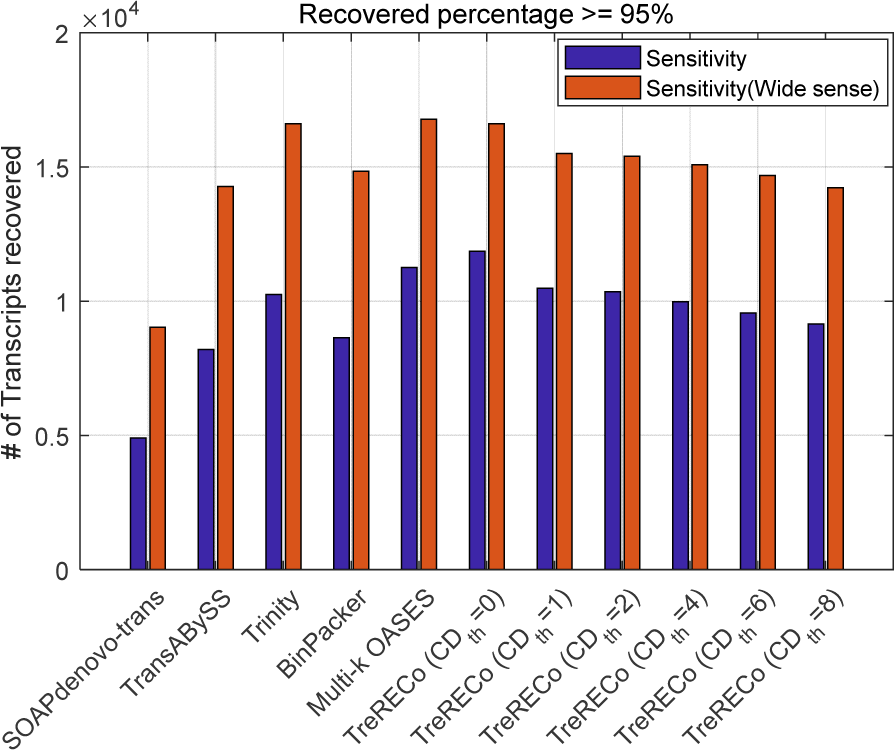
A comparison of (strict sense) sensitivities with wide sense sensitivities for human sample (SRR445718). Target coverage ≥ 95%.

### 2. Results for Simulated Reads

Using simulated reads, we can perform more in-depth investigation on assembler performance, including abundance estimation, as we have prior knowledge on the exact set of isoforms and their expression level, even though the simulated reads may have different characteristics from the real ones and the performance demonstrated for simulated reads could be different from practical performance for real data.

#### Data generation and prior knowledge

The simulated data we used was generated by the Flux Simulator (Griebel *et. al,* 2012) using UCSC mm9 reference genome and its annotation. Flux simulator first randomly generates expression levels for all the transcripts in the annotation and, then simulates the library preparation, including reverse transcription, fragmentation and size selection, to finally obtain reads through sequencing process. The simulator provides various error models, where we used the model for read length 76 and generated 41M reads of length 100. In addition to reads itself, the Flux Simulator also provide the following additional information for each isoform generated, which can be used for in-depth investigation provided in this section.

1. Expressed coverage (covered fraction): the expressed coverage is the percentage of an isoform’s length that is covered by generated reads.
2. Sequenced number: This is the number of reads sequenced for given transcript such that the coverage depth (expression level per base) can be obtain as Sequenced number times read length divided by transcript length times expressed coverage.

#### Parameter setting and pre/post processing

Through a similar procedure for real reads, we compared TraRECo with other assemblers used in the previous sub-section. We used the same parameters for all the assemblers, except that we used *k*-mer length of 31 for Bridger, as it was suggested to use 31 for mouse in its original paper (Chang *et al.*, 2015). Using the candidate isoforms obtained from each assembler, we ran BLASTN to get how many reference transcripts are matched to candidates. Here, we did not use mm9 Ensembl transcriptome. Instead, we used the gene annotation and the expression level profile obtained from the additional information to build the reference transcriptome that contains the exact set of transcripts from which the simulated reads come.

#### Sensitivity versus precision

Table 3 shows the number of all transcript candidates found with each assembler and the number of recovered references with coverage greater than or equal to the specified target value, i.e., 95%, 90% and 80%, respectively. In the last row, we also showed the number of transcripts with its expressed coverage greater than or equal to the specified target value. As a matter of fact, the number of recovered transcripts for given target coverage cannot exceed this number for the same coverage values. The sensitivity can now be defined as the percentage of the recovered transcripts among all reference transcripts with its expressed coverage greater than or equal to the target value. The sensitivity versus precision for simulated reads were shown in Fig.7, where, compared with other assemblers, TraRECo showed the best performance in all assemblers in both sensitivity and precision. Different from real reads, the performance difference between TraRECo and other assemblers are considerable, even though this does not necessarily mean similar performance for real data since the characteristics can be different.

**Table 3.** A comparison of the number of transcripts found and matched to a reference for the simulated reads (Mouse). Target coverage = 95%, 90% and 80%. For TraRECo, the results with *D*_*th*_ =0.05 and *CD*_*th*_ =0, 1, 2, 4, 6, 8, 12 and 16 are shown.

| Assembler | # of<br>candidates<br>found | # of transcripts matched<br>for Target coverage of |  |  |
| --- | --- | --- | --- | --- |
| | | $\geq 95\%$ | $\geq 90\%$ | $\geq 80\%$ |
| SOAPdenovo-trans | 21640 | 4,792 | 5,730 | 6,353 |
| Trans-ABYSS | 29,642 | 5,330 | 6,280 | 6,934 |
| Trinity | 27,962 | 5,499 | 6,314 | 7,067 |
| BinPacker | 19,489 | 5,117 | 5,920 | 6,655 |
| Muli-k OASES | 56,885 | 7,201 | 8,132 | 8,727 |
| Bridger | 22,737 | 6,015 | 6,781 | 7,378 |
| TraRECo, $D_{th}=0.05$, $CD_{th}=0$ | 42,275 | 7,923 | 8,739 | 9,205 |
| TraRECo, $D_{th}=0.05$, $CD_{th}=1$ | 27,005 | 7,700 | 8,511 | 8,975 |
| TraRECo, $D_{th}=0.05$, $CD_{th}=2$ | 26,398 | 7,677 | 8,481 | 8,946 |
| TraRECo, $D_{th}=0.05$, $CD_{th}=4$ | 19,813 | 7,613 | 8,395 | 8,840 |
| TraRECo, $D_{th}=0.05$, $CD_{th}=6$ | 15,540 | 7,540 | 8,298 | 8,704 |
| TraRECo, $D_{th}=0.05$, $CD_{th}=8$ | 13,502 | 7,420 | 8,110 | 8,451 |
| TraRECo, $D_{th}=0.05$, $CD_{th}=12$ | 11,503 | 6,941 | 7,473 | 7,696 |
| TraRECo, $D_{th}=0.05$, $CD_{th}=16$ | 10,290 | 6,408 | 6,821 | 6,997 |
| The number of Transcripts generated with<br>its expressed coverage ≥ Target coverage | | 9,911 | 11,538 | 12,733 |

**Fig.7.**
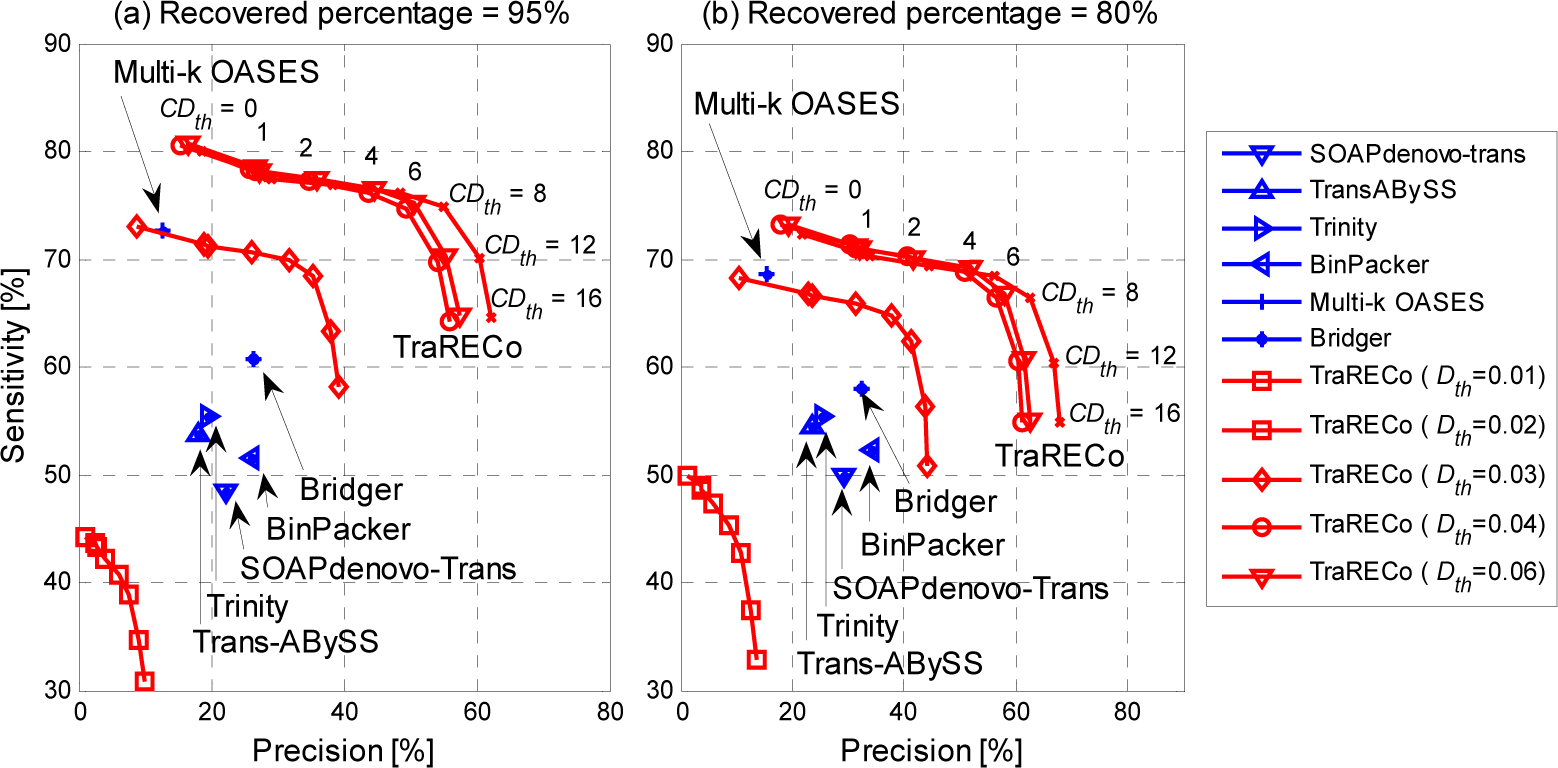
Sensitivity versus precision for the single-ended simulated reads. Target coverage = 95% (a) and 80% (b), respectively. Sensitivity is defined as the number of recovered transcripts divided by the number of reference transcripts with its expressed coverage larger than the target coverage (provided in the last row of Table 3).

#### Transcript length unbalances and wide sense sensitivity

As did for real data, we also checked length unbalances and wide sense sensitivity for simulated data, which are shown in Fig.8 and Fig.9. In Fig.8, we showed only those for Trans-ABySS, multi-*k* OASES and TraRECo (*D*_*th*_ = 0.05, *CD*_*th*_ = 4). Compared with those for real data, the simulated read showed much better balances between the candidate and the reference transcripts and almost no small RNAs were detected. The R^2^ measure for SOAPdenovo-trans, Trans-ABySS, Trinity, BinPacker, multi-*k* OASES, Bridger and TraRECo (*D*_*th*_ = 5, *CD*_*th*_ = 4) were 0.990 (best), 0.962, 0.960, 0.894, 0.732 (worst), 0.944 and 0.961, respectively. Fig.9 shows a comparison of (strict sense) sensitivities with wide sense sensitivities. As can be inferred from Fig.8, one cannot observe considerable difference between (strict sense) sensitivities and wide sense sensitivities.

**Fig.8.**
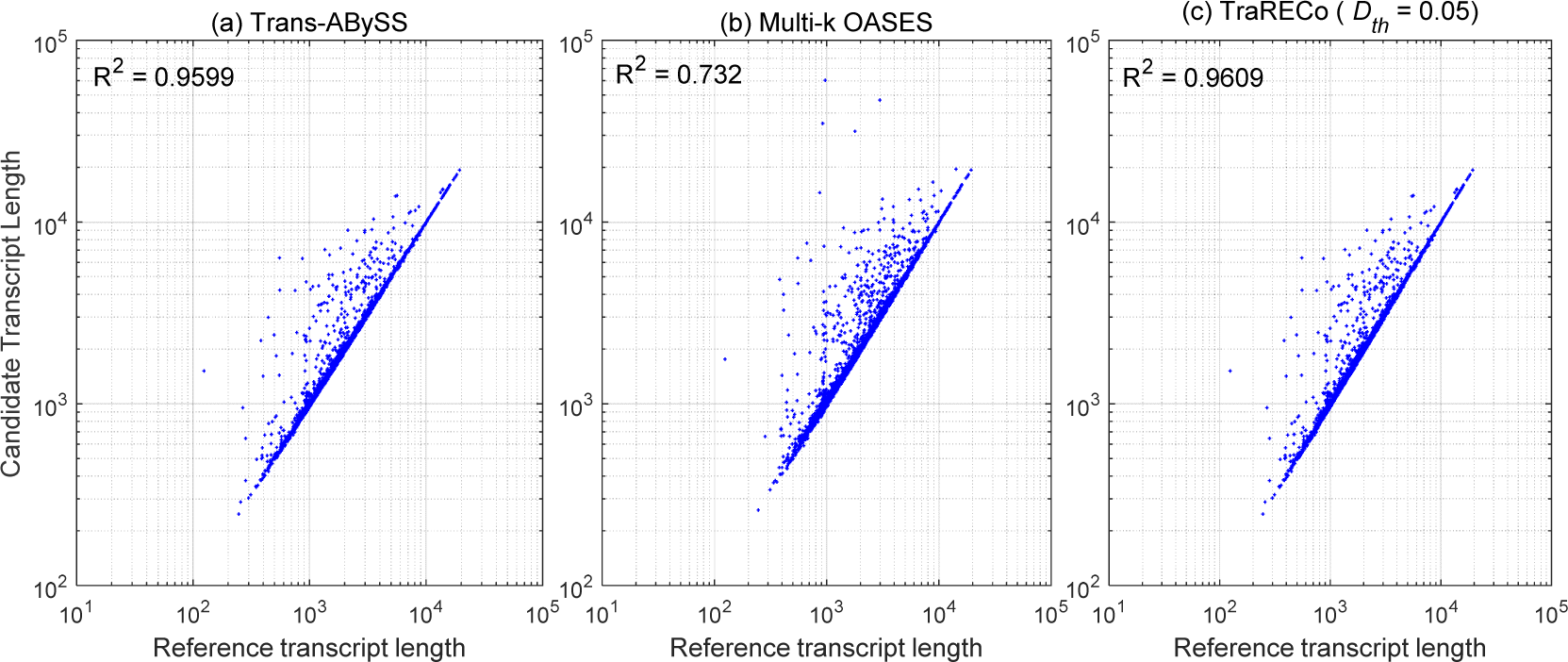
Scatter plot representing length correlation between reference and candidate transcripts matched with 95 or higher coverage of reference for the simulated read. (a) Trans-ABySS, (b) multi-*k* OASES, (c) TraRECo (*D*_*th*_ = 0.05).

**Fig.9.**
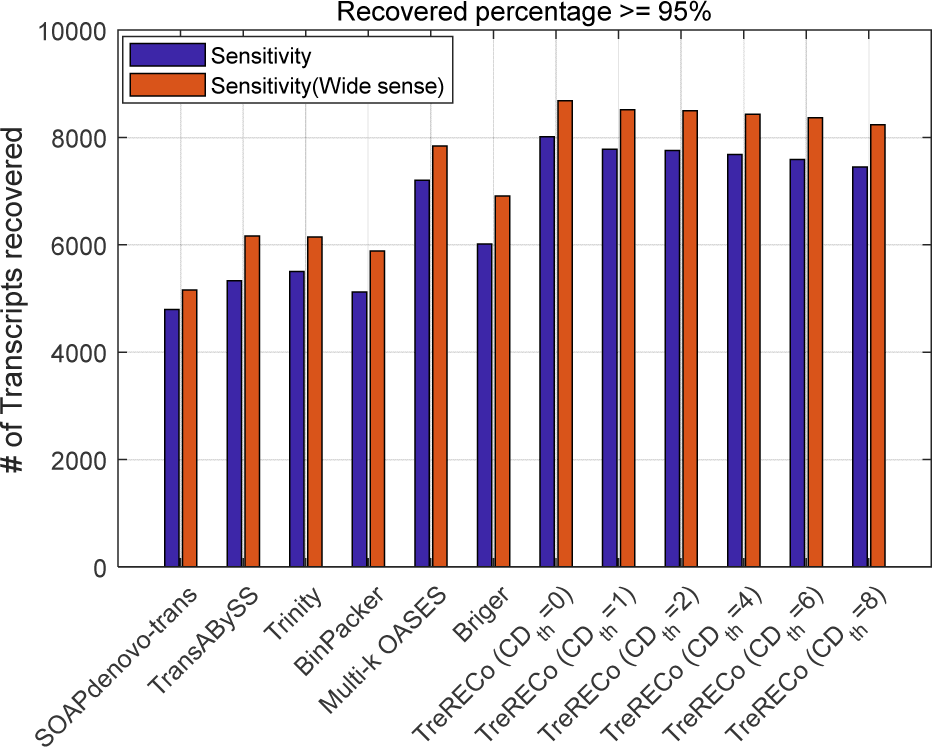
A comparison of (strict sense) sensitivities with wide sense sensitivities for simulated reads. Target coverage ≥ 95%.

#### Abundance and Sensitivity

Fig.10 show the number of undetected transcripts with their true abundances approximately equal to abscissa, where, to give insight into the percentage of undetected transcript, the number of all reference transcripts were also shown. We used 30 bins for true abundance from 0 to 4.2 in log-scale. Fig.10 (a) shows only TraRECo with *D*_*th*_ = 0.01, 0.02, 0.03 and 0.05, which clearly shows the improvement with larger value of *D*_*th*_. Although our expectation was some improvements especially for lowly expressed transcripts, the results shows marginal improvement for lowly expressed transcripts while considerable improvement for transcripts with medium expression level from 20 to several hundred in normal scale (1.3~2.5 in log-scale). When comparing with other assemblers, TraRECo showed slight improvements for transcripts with low expression level. There are two things to note. (1) Most of the assemblers, including TraRECo, however, could not successfully detect those transcripts with their abundance less than 10 (1 in log-scale). (2) Even with medium to high expression level, non-negligible portion of transcripts could not be detected. This is true even if we consider (strict sense) sense sensitivity as it is slightly lower than the wide sense sensitivity as shown in Fig.9. Fig. 11 shows how many transcripts were common among TraRECo, multi-*k* OASES and Bridger for target coverage of 80% or higher. (We selected the three assemblers as they showed highest sensitivity.) In this figure, we see that (1) considerable percentage of candidates (68 to 81%) are common among these assemblers and (2) non-negligible percentage (1,559/11,764 ≈ 13% for abundance ≥ 5 and 251/8,133 ≈ 3% for abundance ≥ 20) still could not be recovered by all these three.

**Fig.10.**
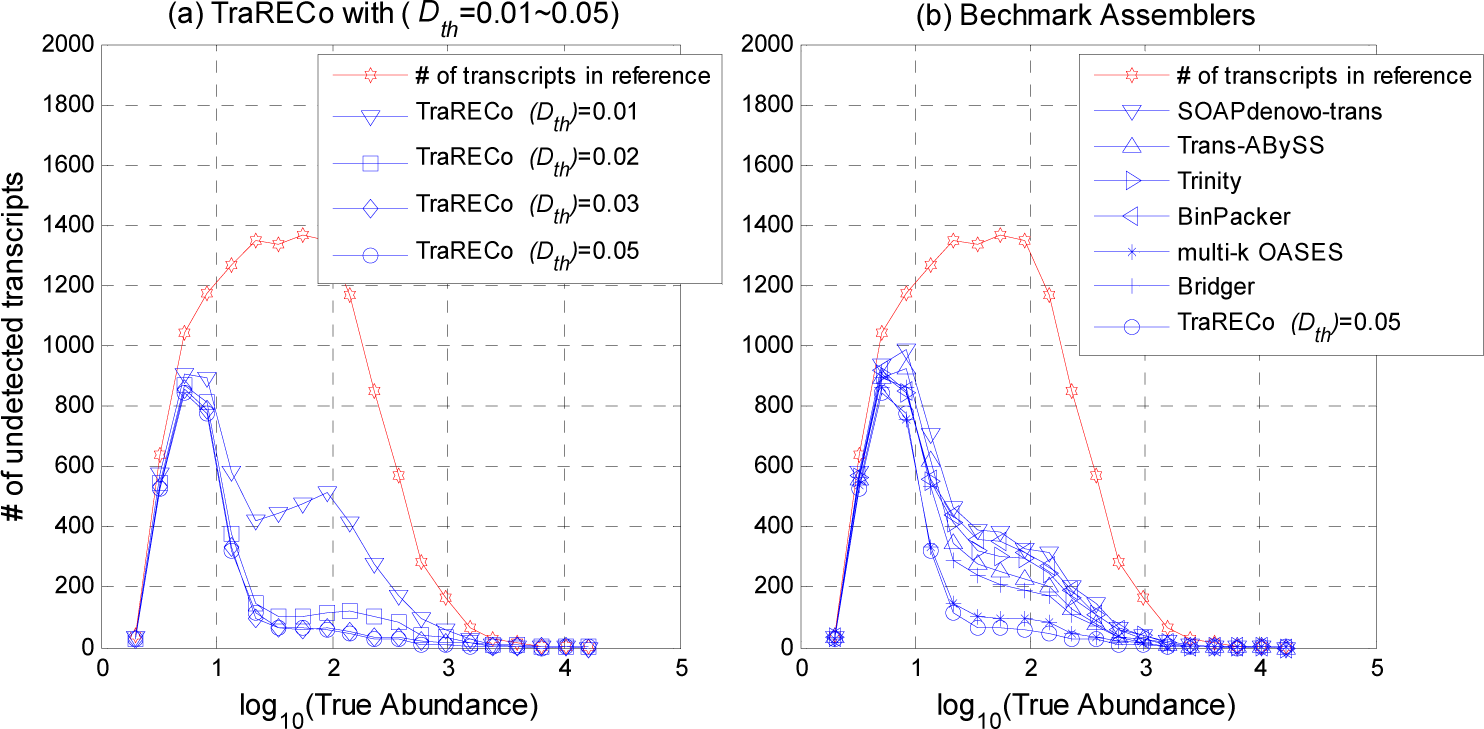
Histogram of the number of undetected transcripts (in wide sense) for simulated reads. Target coverage ≥ 80%. True abundance is shown in log-scale. To give insight into the percentage of undetected transcript, the number of all reference transcripts were also shown.

**Fig.11.**
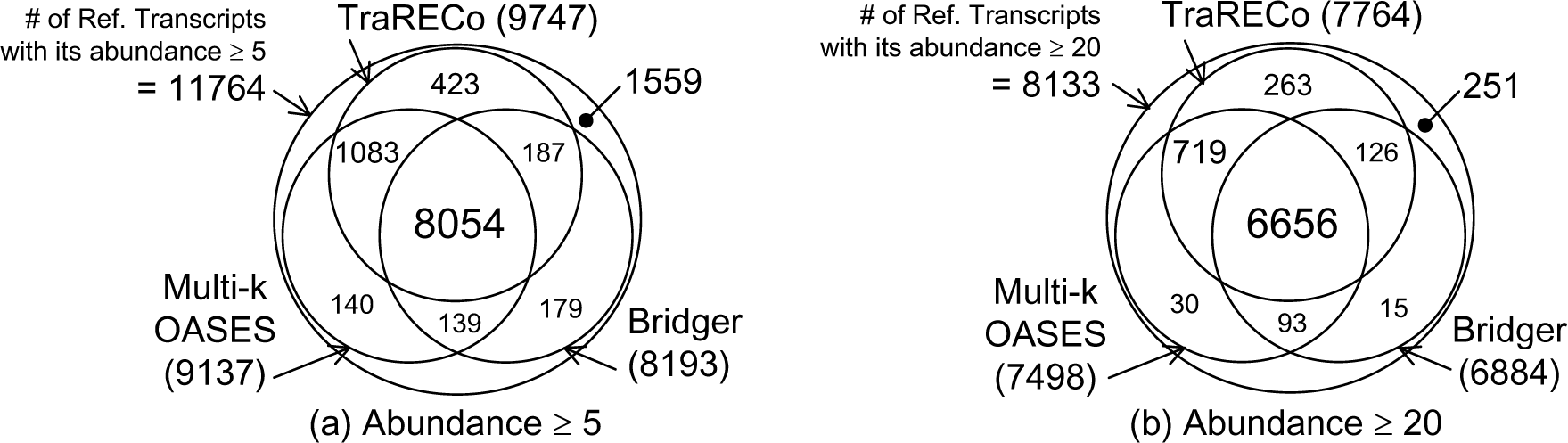
Venn diagram showing the number of transcripts detected that are common among various assemblers for the simulated reads. Target coverage 80%. Considered only those reference transcripts with its abundance being equal to or greater than (a) 5 and (b) 20.

#### Abundance estimation performance

Note that TraRECo provides also the abundance estimates for each candidate, with which we could trade sensitivity with precision by controlling the coverage depth threshold, *CD*_*th*_. Although the abundance estimation is a secondary issue in *de novo* transcriptome assembler, it would be interesting to see how accurate the abundance estimation of TraRECo is. Fig.12 shows a comparison of abundance estimation performance between Trinity+RSEM and TraRECo. The latter produces abundance estimates for all the detected isoforms along the assembly procedure, while Trinity provides a separate abundance estimation package, called RSEM, with which the estimates are obtained by realigning reads to the assembled isoforms. The RSEM provides abundances in Transcripts Per Kilobase Million (TPM) and Fragments Per Kilobase Million (FPKM), while TraRECo provides it in Reads Per Kilobase Million (RPKM) and raw coverage depth. So, we used the effective transcript length and expected read count provided by RSEM to obtain the estimates in RPKM as 10^9^*n*_*k*_/*l*_*k*_*N* where *n*_*k*_ and *l*_*k*_ are respectively the expected read count and effective length of the *k*^th^ transcript and *N* is the sum of *n_k_*’s for all *k*. We selected those isoforms with the coverage of 80% or higher and compared their abundance estimates with the true abundances provided as a prior knowledge. The R^2^ measure for the TraRECo and Trinity+RSEM were 0.684 and 0.728, respectively. Although TraRECo showed less accurate abundance estimates than that of Trinity+RSEM, they are quite close. Another thing to note is that, from Fig.12 (a), TraRECo tends to slightly underestimate the abundances for a large portion of transcripts even though big differences as in Trinity+RSEM are seldom. The under-estimation tendency is because some of the reads were discarded in the contig growing step of TraRECo as they could not be aligned within the specified distance margin.

**Fig.12.**
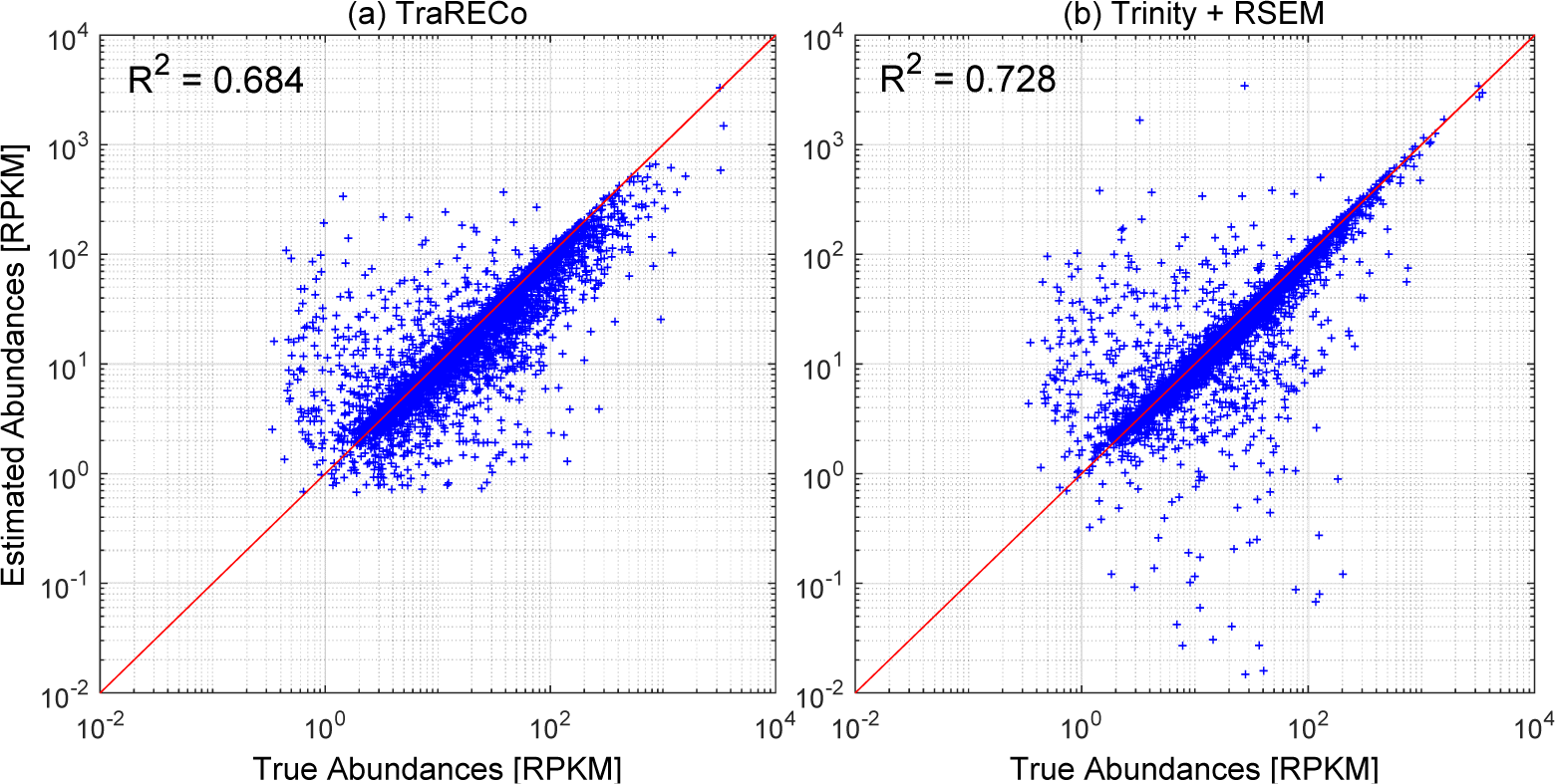
Abundance estimation accuracy of TraRECo (a) and Trinity + RSEM (b)

### 3. Assembly Quality measurement using DETONATE

For real data, we do not have the exact set of transcripts and the evaluation can only be based on known transcripts disregarding the unknown, yet possibly existing, isoforms, while simulated data may have different characteristics from real ones. Given that ground truth unknown for real data, DETONATE (Li et al., 2014) or TransRate (Smith-Unna *et al.*, 2015) can be used for a more reliable measure of *de novo* transcriptome assembly. To provide insight into how well the assembled candidates represents the data in an efficient way, we used DETONATE, which provides two types of assembly quality measure, i.e., RSEM-EVAL without reference transcriptome and REF-EVAL with reference, where we used the former, which shows how well and efficiently the assembled transcripts represent the read data. Table 4 shows the results with RSEM-EVAL method. For real human sample (SRR445718) and the simulated reads, the best RSEM-EVAL scores were −1,391,489,463 and −1,480,014,038, respectively, both with TraRECo (*CD*_*th*_ = 2) while the best scores among other assemblers were −1,450,049,444 with TransABySS and −1,482,875,599 with Trinity, respectively. For real mouse sample, the best score of TraRECo was −11,282,292,195 with *CD*_*th*_ = 16, while the best among other assemblers was - 11,260,544,990 with SOAPdenovo-Trans. For the real mouse sample (SRX062280), it seems that the number of candidate transcripts was the dominant factor to get a better score as TraRECo with *CD*_*th*_ = 16 (having the least number of candidates) obtained the best among all TraRECo scores and gradually got worse with smaller *CD*_*th*_.

**Table 4.**
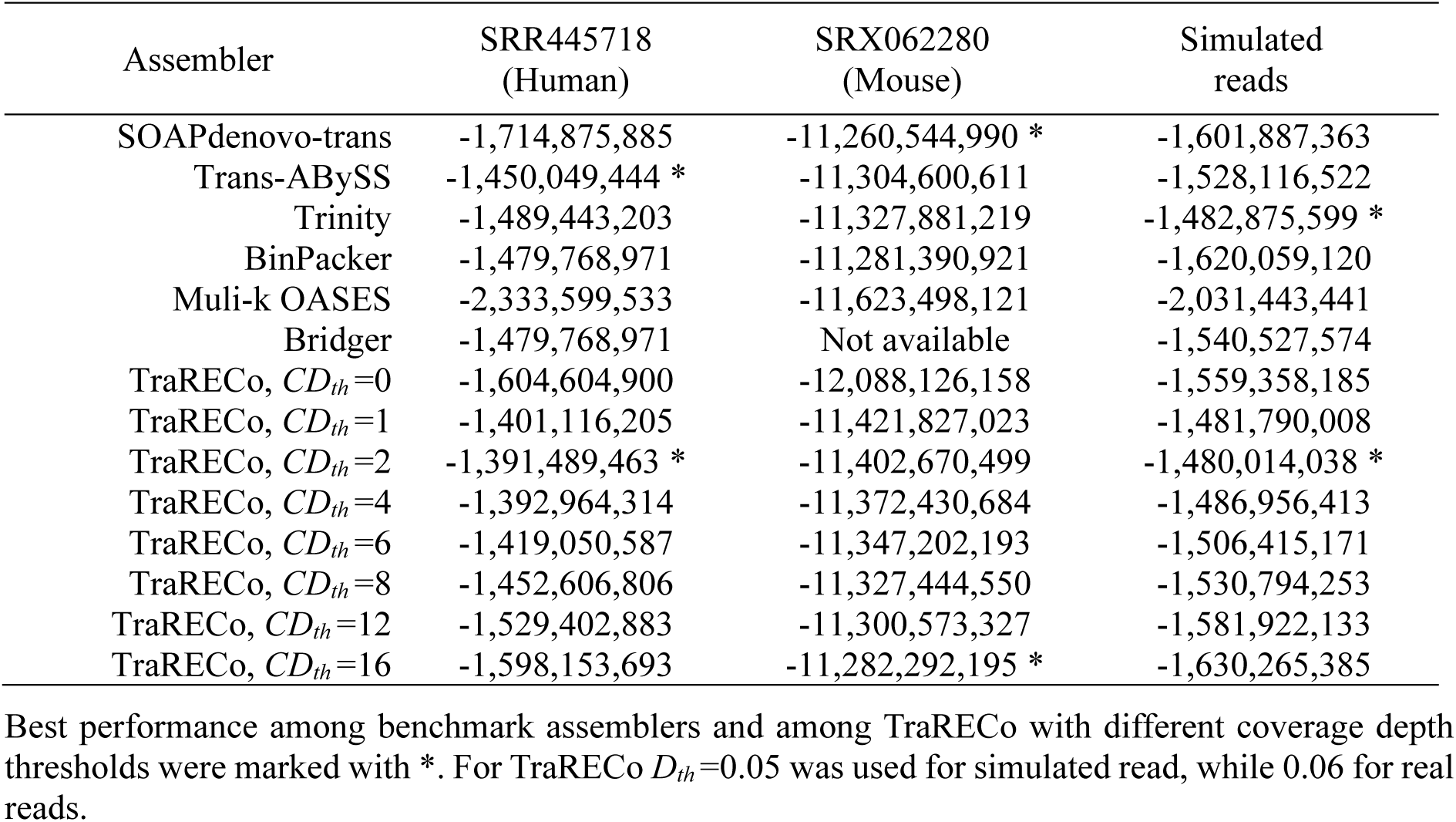
A comparison of DETONATE (RSEM-EVAL) scores for the three data samples.

### 4. Computational Burden and Run Time

Most of the existing assemblers took around 2 to 20 hours, except for multi-*k* OASES which ran a few days to perform Velvet many times for different *k*-mers and combining them to obtain final splicing graph. Comparing run time with the existing ones, TraRECo took around 64 hours for the simulated reads and 150~180 hours for real reads with 16GByte memory, which are much longer than those with existing assemblers. There are two reasons for the much longer running time of TraRECo. First, TraRECo is using direct alignment of reads to contigs, which has a computational complexity grows, in the worst case, as (*NL*)^2^, where *N* is the number of reads and *L* is the read length. This is the bottleneck of the running time in TraRECo. One thing to note is that, as the contig growing step compares reads with contigs, not with other reads, the complexity grows as *NL*⋅*N*_*c*_*L*_*c*_, where *N*_*c*_ is the number of contigs and *L*_*c*_ is the average contig length. Since *N*_*c*_*L*_*c*_ is typically much smaller than *NL*, the computational complexity might be far less than what is expected with a square growth. A rough estimation of *N*_*c*_*L*_*c*_ is *NL* divided by the average coverage depth (abundance), which, as we saw in Fig.8, is approximately one hundred. Another reason is that TraRECo was developed using MATLAB™, a proprietary software tool, to reduce debugging time, while most of assemblers were using C/C++. As MATLAB™ is typically much slower than those developed using C/C++, we can save some proportion of running time if we use C/C++, which is a software development issue rather than a bioinformatics issue. Most of all, as our focus is on the development of new methods, not on software development, the running time was considered as a secondary issue in this study.

## Discussion

Sensitivity and precision are two primary performance measures of *de novo* transcriptome assemblers and there certainly exists tradeoff between the two criteria. To maximize these performances, many *de novo* transcriptome assemblers perform assembly in two steps: (1) building splicing graph and (2) searching for plausible paths, where, for the former step, de Briujn graph approach was widely adopted by most of the existing assemblers since its computational burden is only linearly increasing with the read data size. One problem in de Bruijn graph is that it is not well suited to combat read errors and sequence repeat. To overcome this problem Schulz *et al.*, (2012) proposed a multi-*k* approach, where de Bruijn graphs are constructed separately for many different *k* values. Although it took much more times to obtain the final splicing graph, multi-*k* OASES provided the highest sensitivities among all existing assemblers for all the samples including simulated read. In precision, however, multi-*k* OASES was shown to be the worst among all assemblers, while the best was the recently proposed BinPacker, especially for human sample. In between these two extreme cases, Bridger and Trinity were shown to well compromise the two performance criteria.

Compared to these de Bruijn graph approaches, TraRECo provides a new framework for *de novo* transcriptome assembly by combining the consensus matrix-based error correction procedure with the direct read alignment based on greedy approach. Through the work presented in this study, we could confirm that the proposed contig growing procedure using consensus matrices can combat read errors efficiently in the sense that the sensitivity of TraRECo with low coverage depth threshold (*CD*_*th*_) were shown to even better than the multi-*k* OASES. This improvement could be achieved by making more erroneous reads to be participated in the graph construction step, which, in turn, improve the quality of read depth information used for the subsequent steps, i.e., searching for plausible paths. This aspect is certainly different from the simple read-error removal or error correction based on topological structure of the de Bruijn graph and the difference can make the proposed approach an alternative method, at least, for splicing graph construction step, even though it has a bottleneck of square complexity with read data size.

On the other hand, the direct alignment of reads to build contig made us able to resort full connection information of short read to suppress the impact of sequence repeat of length less than the minimal overlap width used for read alignment. The improvements over a broad range of abundances shown in Fig.10 for simulated reads support this arguments. Although there is still problem that transcripts with low expression level could not be connected during contig growing step, we could alleviate the problem by connecting contigs with smaller connection threshold *C*_*th*_ before performing junction search and graph construction step. Resorting full connection information of a read played also an important role in junction search and group (graph) decomposition step as well, where we could utilize the junction overlap width for group/graph decomposition to facilitate the subsequent joint isoform detection and abundance estimation.

At the final step, we borrowed the idea of IsoLasso to detect isoforms and estimate its abundance jointly. IsoLasso is a simple, yet quite powerful, method for the subsequent path search step, even though there are more sophisticated approaches, such as Bridger and BinPacker. Our approach, however, had difference from IsoLasso in that we allow the number of inclusion of a segment participating in a path can be more than one by taking account loopy graphs, which never occurs in reference-based assembly as they utilize known gene annotation, while they can be appeared in *de novo* assembly as an artifact of sequence repeat.

The results from the simulated reads also showed that many challenges are still remaining to be solved, especially for path search step, since a non-negligible portion of transcripts with their expression level being high enough (read depth greater than 20) still could not be recovered by existing *de novo* assemblers, including TraRECo. This seemed to be due mainly to sequence repeats by which many transcripts/isoforms were merged together and one may need to devise more sophisticated methods to decouple merged transcripts/isoforms and artificial gene fusion.

## Conclusions

Many existing *de novo* transcriptome assemblers are based on the de Bruijn graph, which builds splicing graph in linear time of data size while suffers from read errors that make the splicing graph complicated. Based on this point of view, it looks natural that the recent works, such as Bridger and BinPacker, were focusing more on reliable path search to improve precision. Another research direction was made by (Schulz *et al.*, 2012) to suppress the impact of read errors and short repeat by using multiple *k*-mers approach. The study presented here pursued the same objective as the multiple *k*-mers approach and we believe it was successful in the sense that the proposed approach showed the highest sensitivity if we do not consider precision. TraRECo showed also a good performance even when comparing both sensitivity and precision at the same time. Although the computational burden of direct read alignment can be much higher than the single *k*-mer de Bruijn graph approach, it seems not too bad as its computational burden is far less than the worst square complexity due to its recursive computation providing us a potential alternative as a benchmark. Overall, TraRECo could provide a reliable splicing graph construction, which is an important issue since *de novo* assembly is mainly to explore not-yet-discovered isoforms and must be able to represent as much reads as possible in an efficient way.

## Methods

The entire procedure of the proposed assembly consists of three parts: (1) contig growing, (2) junction search and graph construction, and (3) joint isoform detection and abundance estimation as briefly shown in Fig.13. The contig growing utilizes greedy approaches widely adopted for DNA assembly, e.g., (Zhang *et al.*, 2004), SSAKE (Warren *et al.*, 2007) SHARCGS (Dohm *et al.*, 2007) and VCAKE (Jack *et al.*, 2007), as it is more suitable for error correction and one can utilize full connection information of a read. The second step is to search junctions among contigs and then to construct graph, which consists of nodes (representing a segment of base sequence) and edges (representing the connection between segments). In these steps, the read coverage for each base is tracked to obtain the coverage depth profile for each contig and segment. Finally, these information are used to detect isoforms and estimate abundances jointly. This procedure is distinguished from previous methods as follows:

**Fig.13.**
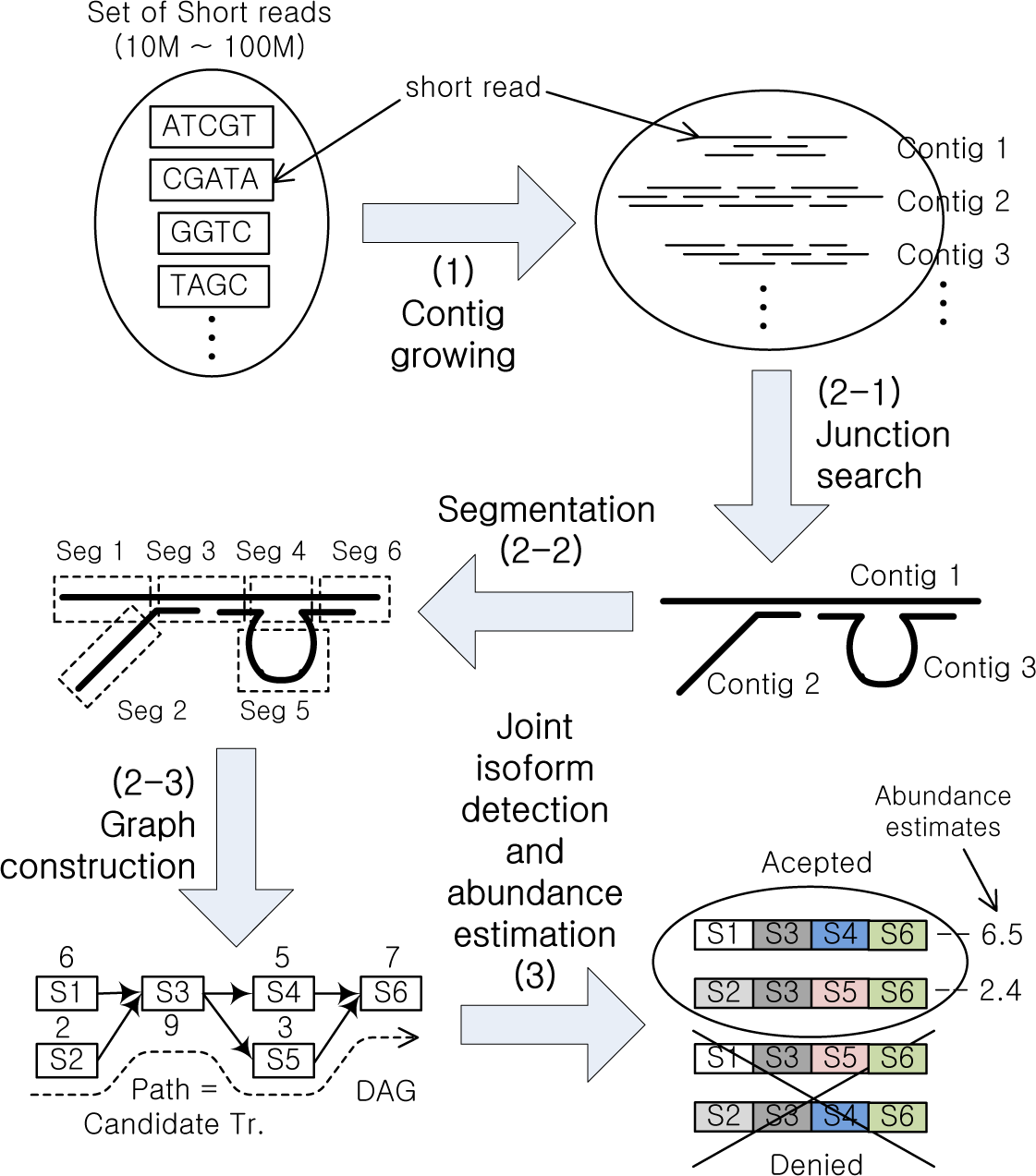
The entire workflow of TraRECo assembly procedure consisting of (1) contig growing, (2) junction search and splice graph construction and (3) joint isoform detection and abundance estimation.

1. In contig growing step, we use “consensus” matrix, which holds alignment profile represented by base count for each location of a contig. This profile made it possible to identify errors and check if any similar sequences merged into a single contig.
2. This alignment profile was also tracked in the subsequent junction search and graph construction step to be delivered to the final stage, where one can jointly detect isoforms and estimate abundances.

Throughout this section, we will provides some more details of the assembly process, highlighting the key features of the proposed assembler.

### 1. Contig growing based on consensus matrix

Contig grower builds contigs by aligning short reads to all the contigs in the contig pool in recursive manner. It proceeds as follows: Let Ψ be the set of contigs found so far and Ξ be the set of short reads. We selected a read from Ξ and try to align it with all the contigs in Ψ. Then, we chose the contig that had the widest overlap with the read selected. If the read completely overlapped with the contig, we merged the read to the contig, while we extended the contig if the read only partially overlaps. If there was no contig having overlap longer than or equal to a predefined value, we simply added the read to Ψ as a new contig. This procedure was repeated until all the reads in S are processed. One key feature in our contig growing was that we used consensus matrix to provide read error correction, as briefly shown in Fig.14.

**Fig.14.**
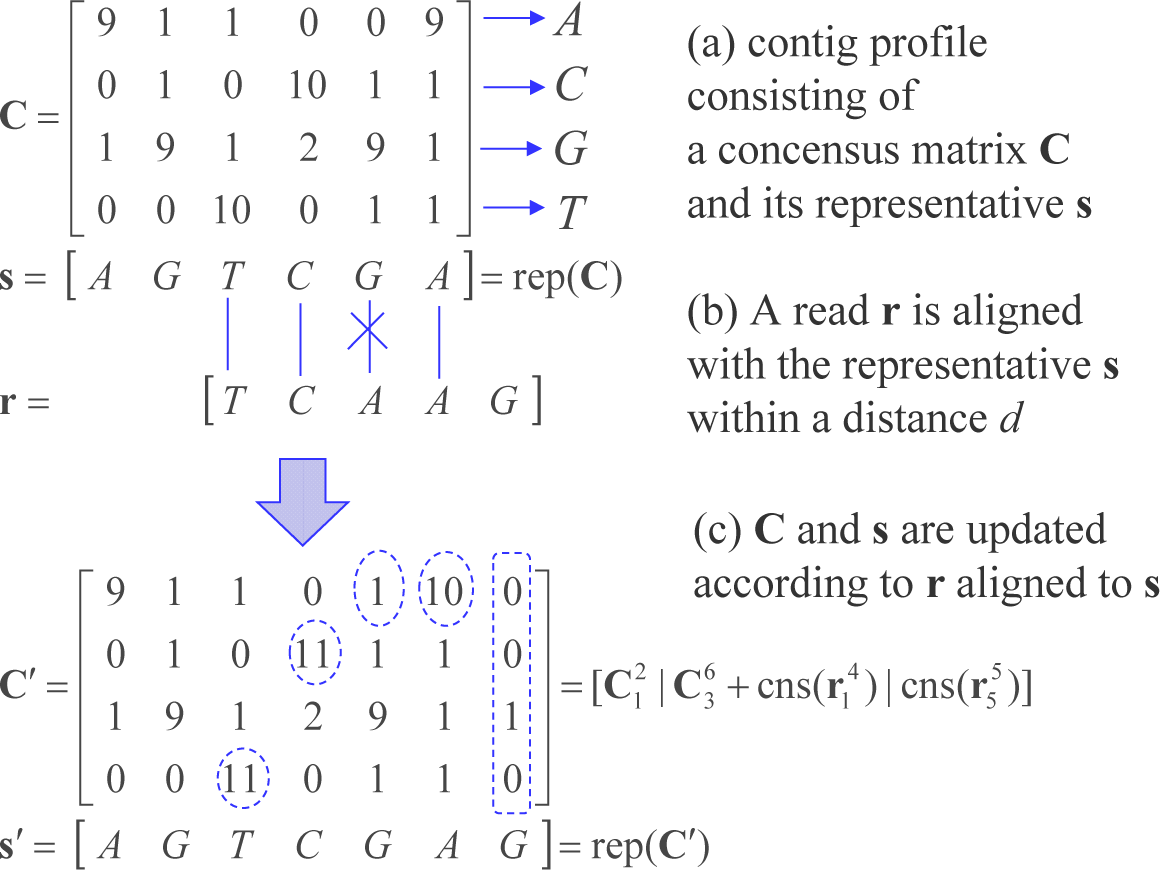
An example of contig profile update when a read **r** of length 5 partially overlaps with **s** = rep(**C**) of length 6 from the right (differing only one base among 4). The contig profile **C** is extended to and replaced with the updated contig profile **C**′, which is now of length 7. The updated elements are marked with dashed circle.

In the proposed scheme, a contig of length *l* is represented by a consensus matrix, **C** of size 4x*l*, and the corresponding representative vector **s** of length *l*. Each row of **C** corresponds to a base in {A, C, G, T} and each element of **C** is the base count of reads aligned to this contig. On other hand, each element of s corresponds to the row index with the highest count at each column of **C**, i.e., *s*_*j*_ = arg max,._*i*∈{1 2 3 4}_ *c*_*i,j*_, The representative **s** is used to test alignment with a read as follows: Let us consider alignment test of a read **r** of length *m* with a representative **s** of length *l*. Without loss of generality, we assume *l* ≥ *m*. Three cases can happen: (1) complete overlap, (2) partial overlap from the left or right of the reference, **s**, and (3) no overlap. Let 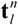 be a portion of a vector **t** such that 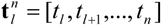 for *l* ≤*n* and 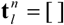, a null vector of length 0, if *l* > *n*. The same notation can also be applied to a matrix, i.e., 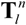 is a matrix consisting of the *l*^*th*^ to *n*^*th*^ column of matrix **T**. Then, one can write 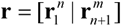 for an integer 0 ≤ *n* ≥*m*. Based on this notation, we say that **r** completely overlaps with s if for some integer *a* in [1, *l m*+1] 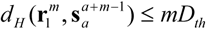 with *m*≥*O*_*th*_, where *d*_*H*_ (**a, b**) is the Hamming distance between the two vector **a** and **b**, *D*_*th*_ is the normalized distance threshold such that *mD*_*th*_ is the number of errors allowed and *O*_*th*_ is the minimum overlap width for the overlap to be considered as valid one. On the other hand, we say that **r** partially overlaps with **s** if there exists *n* satisfying one of the following conditions: (1) 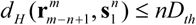 (partial overlap from the left) and (2) 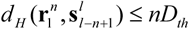 (from the right) for *m* > *n* ≥*O*_*th*_. If there exist multiple *n* satisfying the conditions, we take the largest value. For notational simplicity, we define two functions: rep(⋅) and cns(⋅). The function rep(⋅) takes a consensus matrix **C** and returns its representative sequence such that **s** = rep(**C**), while the function cns (⋅) takes a sequence s and returns a consensus matrix initialized by **s**, i.e., each element of **C** = cns(**s**) is set to *C*_*i,j*_ = 1 for *i* = *S*_*j*_ or 0 otherwise. Now, we can describe the contig profile update procedure using the two functions. We consider 3 cases. Suppose that a read **r** of length *m* partially overlaps with the representative **s** = rep(**C**) of length *l* ≥*m* from the left/right of **s** and the overlap width is *m* ≥*O*_*th*_. Then, we update the contig profile as

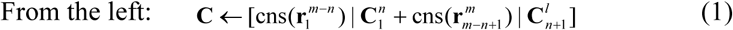

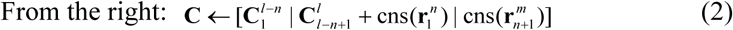

An example of contig profile update is shown in Fig.14 for this partial overlap case. Suppose that the read **r** of length *m* completely overlaps with the representative **s** = rep(**C**) of length *l* ≥*m* at position *a.* Then, we update the contig profile as

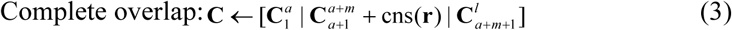

If the read **r** does not overlap with **s** or overlaps with a width less than *O*_*th*_, we simply add cns(**r**) to the contig pool as a new seed. The overall contig growing procedure is as follows.

Contig growing

*Input*: **Ξ** = {**r**_1_,**r**_2_, …,**r**_*N*_}

*Initialization:* Set seed contig, e.g., Ψ = {cns(**r**_1_)} and remove **r**_1_ from Ξ

*Loop*: For all **r** ∈ Ξ

A. Test alignment of **r** with **s** = rep(**C**) for all **C** ∈ Ψ

B. If there exist **C** having overlap width *n* larger than *O*_*th*_ (within the distance *nD*_*th*_), select C having the widest overlap with **r** and update **C** using (1), (2) or (3) based on the overlap width and position

C. Otherwise, add **r** as new contig, i.e., Ψ ← Ψ + cns(**r**)

*End loop*

*Output*: Ψ = {**C**_*k*_: *k*=1,2,…}

#### Parameter setting

Two key parameters in the proposed contig growing procedure are the normalized distance threshold *D*_*th*_ and the overlap threshold *O*_*th*_. *D*_*th*_, is normalized value per base such that *nD*_*th*_ is the maximally allowed number of different letters for a portion of a read of length *n* to be aligned with a contig representative, where *n* ≥*O*_*th*_. If *D*_*th*_ is too small, reads with more errors will not be aligned with a target contig and may eventually be disregarded. On the other hand, if it is too large, more similar sequences from other isoforms can be merged together resulting in artificial gene fusion. Such artifacts make the splice graph complex and the subsequent joint isoform detection and abundance estimation complicated, exactly in the same way as the de Bruijn graph-based approach suffers.

In contrast to de Bruijn graph-based approach, however, the artificial gene fusion from short repeats can be avoided by setting *O*_*th*_ relatively large (larger than the *k*-value in de Bruijn graph-based approach) as long as the read length is much longer than typical *k*. Note also, however, that it is undesirable to set *O*_*th*_ too large, especially when read coverage is not enough, i.e., for isoforms with low expression level. If one set *O*_*th*_ to large, true transcripts cannot eventually be connected on those regions where the read coverage is low. On the other hand, if one set it too small, the graph construction can be vulnerable to short repeats resulting in possible merge of multiple isoforms with similar sequences. As a matter of fact, there is a tradeoff between the error correction capability and the complexity of the final splice graph to be used for joint detection and abundance estimation and we need care to set the thresholds, *D*_*th*_ and *O*_*th*_.

#### Post contig combining and contig filtering

Post contig combining can be helpful especially for those isoforms with low expression level, which can be performed in the same way as contig growing, but with a smaller overlap threshold, for which we defined connection threshold denoted as *C*_*th*_. Although setting *C*_*th*_ smaller than *O*_*th*_ can make undesired contigs connected to each other due to sequence repeat, it can be identified at the junction search stage and be eventually resolved through the subsequent steps. One also can remove those nodes with its length less than a certain threshold and its read coverage depth is less than, for example, 2 since they are highly likely to be short fragments that could not be merged due to many errors.

### 2. Junction Search and Graph Construction

#### Junction search and contig grouping

In the second step of the procedure, we first search junctions among contigs by testing alignment of prefix and suffix of a contig with other contigs. It is exactly the same procedure as the alignment test in the contig growing step. Note that (1) the overlap width around a junction is smaller than the nominal read length as we employed greedy approach and (2) we need to carefully set the junction overlap threshold, say *J*_*th*_, since it plays the same role as the overlap threshold in contig growing and filtering. 3hen, based on the junction information collected for all pairs of contigs, one can group contigs that are linked together, where each group (hopefully) represents a gene with multiple isoforms. In this work, we set *J*_*th*_ = 32.

#### Junction filtering and group decomposition

Sometimes, group size appears to be very large, which corresponds to a complex graph observed in de Bruijn graph-based approaches. In this case, the isoform detection that follow become very complicated. At this stage, one can invalidate some junctions according to their confidence. In this approach, one can utilize not only the read coverage depth of the two contigs involved in each junction, but also the overlap width and distance within the overlap region. As a matter of fact, it would be quite risky to apply a fixed threshold to invalidate junctions because overlap width around junction is roughly proportional to isoform expression level which is quite uneven among genes. Although more sophisticated junction filtering can be devised, we applied a simple filtering as follows: For given junction information of a group, we iteratively invalidate the junction with smallest overlap widths until group size become less than or equal to the predefined number, say 40. See (Peng *et al.*, 2013) for a reasonable group size.

#### Segmentation and graph construction

Now, for each group of contigs, we segment contigs (contig profiles) at every valid junction (overlap) boundaries, and build segment connection matrix to finally construct splice graphs for each group of contigs having valid junctions to each other. A splice graph *G(N,E)* consisting of nodes *N* and edges *E* is a directed graph, possibly with loops due to sequence repeats longer than *J*_*th*_. In the splice graph, each node represents a segment having segment profile (consensus matrix) and each edge connects one segment to another. The read coverage profile **v** of a segment with segment profile **S** of size 4xl is obtained by summing all the rows of **S**, column by column.

#### Compacting graph to make it minimal

Define indeg(n) and outdeg(n) be the number of input and output edge degrees, respectively, of a node *n* and src(*e*) and dest(*e*) be the source and destination of an edge *e*, respectively. We say two nodes *n* = src(*e*) and *n′* = dest(*e*) are singly connected iff outdeg(*n*) = indeg(*n′*) = 1. Without loss of information, one can combine these two nodes together to make the graph minimal where no nodes are singly connected. By performing such combining, we will later on assume that the splice graph to be used for isoform detection and abundance estimation is minimal.

### 3. Joint isoform detection and abundance estimation

With a splice graph *G*(*N*,*E*) consisting of nodes *N* and edges *E*, one can now jointly detect isoforms and estimate their abundances based on the per-segment average coverage depth {*y*_*j*_: *j*∈*N*}. To this end, we tried IsoLasso (W. Li, J. Feng, and T. Jiang, 2011). To make it fit into our framework, we slightly modified the procedures.

Let Π be the set of all maximal paths starting from a node with input degree 0 and end at a node with output degree 0. Typically, the number of all paths |п| is larger than the number of true isoforms and the problem is to find the true isoforms only. At first, one can resort pair-end reads to filter out those paths that are not compatible with any pair-end reads used in the contig growing step. However, even with such a filter, the problem still remains, i.e., there exists many branches caused by read errors, which are all compatible with reads as they came from the same set of reads. In order to select a plausible set of candidate isoforms, taking into account the case when there are too many paths, one can use IsoLasso (W. Li, J. Feng, and T. Jiang, 2011), which was proposed to ease the problem of under-determined system. It is a constrained minimum mean squared error estimator that can be concisely written as a quadratic optimization problem:

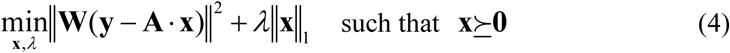

where **x** is |п|x1 vector of which each element is the abundance of a candidate isoform to estimate, **y** is |*N*|x1 vector of which the *i*^*th*^ element is the average read coverage depth of the *i*^th^ node(segment) obtained from the corresponding segment profile, **A** = [*a*_*i,j*_] is |*N*|*x*|п| matrix, where *a*_*i,j*_ is a non-negative integer representing the number of inclusions of the node *i* in the path *j* and **W** = [*w*_*i,j*_] is |*N*|x|*N*| diagonal matrix with 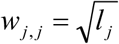, where *l*_*j*_ is the segment length corresponding to the *j*^*th*^ node. **x**≿**0** represents the constraint that all the elements of **x** must be non-negative. We used a variable step gradient search algorithm to find **x** for a set of values of *μ* to minimize (4), after which we took those paths (candidate isoforms) with its length larger than a length threshold, *L*_*th*_, and its estimated abundance larger than a coverage depth threshold, *CD*_*th*_. IsoLasso is a good option especially when |п| ≫ |*N*|, in which case one can reduce the support set by increasing parameter, *μ*.

Final step is to discard those candidates with their length shorter than a length threshold, *L*_*th*_, and their abundance less than a coverage depth threshold, *CD*_*th*_. A typical value of *L*_*th*_ is 200 if we do not take short non-coding RNAs into account. On the other hand, setting *CD*_*th*_ needs cares. With high *CDth*, one can obtain a better precision while true isoforms with low expression level can be removed so that sensitivity will be degraded, and *vice versa.* Although a better joint detector/estimator can be designed to further improve performance, we use this rather simple estimator as our focus is on the proposed contig growing and graph construction scheme demonstrated in previous subsections.

#### Dealing with loopy graphs

As mentioned, each element of **A**, *a*_*i,j,*_ can be larger than 1, which means that we allow a node can be included more than once to resolve loopy graphs, which is caused by relatively long sequence repeats. Although it occurs seldom, we allowed one node in a loopy graph can be included twice, i.e., by assuming a sequence repeat can be occurred only once, we discard those paths that have any nodes included more than twice or more than two nodes included more than once.

## Declarations

### Ethics approval and consent to participate

Not applicable.

### Consent for publication

Not applicable.

### Availability of data and materials

TraRECo and its compiled version (binary executable for windows) are available at Sourceforge, https://sourceforge.net/proiects/trareco/(Files menu). Simulated reads and assembly results have been deposited to Figshare, http://figshare.com, and are accessible with the keyword ‘TraRECo’. The available data include

1. The simulated read and the additional files generated by the Flux simulator.
2. Assembled transcriptomes generated by all the assemblers considered in this paper.
3. TraRECo workshops for the 3 data sets, including reads, reference transcriptome and configuration file that were used to generate the results for TraRECo.

### Authors’contributions

S. Yoon, K. Kang and W.J. Park developed the key idea. S. Yoon wrote code for TraRECo. D. Kim and S. Yoon ran assemblers and performed all experiments. All authors wrote, read, and approved the final manuscript.

